# An annotated consensus genetic map for *Pinus taeda* L. and extent of linkage disequilibrium in three genotype-phenotype discovery populations

**DOI:** 10.1101/012625

**Authors:** Jared W. Westbrook, Vikram E. Chhatre, Le-Shin Wu, Srikar Chamala, Leandro Gomide Neves, Patricio Muñoz, Pedro J. Martínez-García, David B. Neale, Matias Kirst, Keithanne Mockaitis, C. Dana Nelson, Gary F. Peter, John M. Davis, Craig S. Echt

## Abstract

A consensus genetic map for *Pinus taeda* (loblolly pine) was constructed by merging three previously published maps with a map from a pseudo-backcross between *P. taeda* and *P. elliottii* (slash pine). The consensus map positioned 4981 markers via genotyping of 1251 individuals from four pedigrees. It is the densest linkage map for a conifer to date. Average marker spacing was 0.48 centiMorgans and total map length was 2372 centiMorgans. Functional predictions for 4762 markers for expressed sequence tags were improved by alignment to full-length *P. taeda* transcripts. Alignments to the *P. taeda* genome mapped 4225 scaffold sequences onto linkage groups. The consensus genetic map was used to compare the extent of genome-wide linkage disequilibrium in an association population of distantly related *P. taeda* individuals (ADEPT2), a multiple-family pedigree used for genomic selection studies (CCLONES), and a full-sib quantitative trait locus mapping population (BC1). Weak linkage disequilibrium was observed in CCLONES and ADEPT2. Average squared correlations, R^2^, between genotypes at SNPs less than one centiMorgan apart was less than 0.05 in both populations and R^2^ did not decay substantially with genetic distance. By contrast, strong and extended linkage disequilibrium was observed among BC1 full-sibs where average R^2^ decayed from 0.8 to less than 0.1 over 53 centiMorgans. The consensus map and analysis of linkage disequilibrium establish a foundation for comparative association and quantitative trait locus mapping between genotype-phenotype discovery populations.

## INTRODUCTION

Over one billion *Pinus taeda* L. (loblolly pine) seedlings are planted each year in the U.S. in 13 million hectares of plantations that extend from Eastern Texas to Delaware (Mckeand *et al.*, 2003; Smith *et al.*, 2007). Southern pine plantations, composed primarily of *P. taeda* and *P. elliottii* (slash pine), supply 60% of wood products in the United States and 18% worldwide (Prestemon & Abt, 2002). Six-fold gains in stem volume per hectare have been made over the last 75 years through intensification of silviculture and selective breeding (Fox *et al.*, 2007).

Genomics is revolutionizing basic and applied research in this economically important conifer species. Genomic selection, which aims to predict breeding values from the summed effects of genome-wide genetic markers (Meuwissen *et al.*, 2001), has the potential to accelerate the current breeding cycle of *P. taeda* from 12-20 years to less than seven (Resende *et al.*, 2012). Numerous association genetic studies in *P. taeda* have detected significant SNP variants underlying fungal disease resistance (Quesada *et al.*, 2010; Quesada *et al.*, 2014), wood properties (Gonzàlez-Martinez *et al.*, 2007; Chhatre *et al.*, 2013), stem terpenes and other metabolites (Eckert *et al.*, 2010b; Westbrook *et al.*, 2013; Westbrook *et al.*, 2014), physiology (Cumbie *et al.*, 2011), geographical adaptation (Eckert *et al.*, 2010a,b), and gene expression (Palle *et al.*, 2013). A draft assembly of the 22 Gb *P. taeda* genome, the largest genome sequenced to date, was recently reported (Neale *et al.*, 2014; Wegrzyn *et al.*, 2014; Zimin *et al.*, 2014).

A high-density consensus linkage map that is primarily based upon polymorphisms within genes will be useful for all areas of genomic research in *P. taeda* and other conifers. A consensus genetic map will be useful for estimating variation in recombination rates along chromosomes when a physical map for *P. taeda* becomes available (Chen *et al.*, 2002). For genomic selection, a high-density genetic map may be used to design low-density panels of markers that reduce genotyping costs without sacrificing prediction accuracy (Habier *et al.*, 2009). In addition, a consensus genetic map can be used to compare the locations of genetic associations and quantitative trait loci (QTL) discovered in independent populations (Westbrook *et al.*, 2014).

A consensus map may also be used to compare the extent of linkage disequilibrium (LD) within genotype-phenotype discovery populations. Linkage disequilibrium or the non-independence of segregating alleles at different genomic loci may arise from two loci being in close proximity on a chromosome, thus reducing the probability of recombination between them (Flint-Garcia *et al.*, 2003). Statistical power to detect an association between a marker and a trait is inversely proportional to the squared correlations (R^2^) between alleles at a marker locus and the causal variant (Pritchard & Przeworski, 2001). Thus, quantifying the extent of LD is useful for knowing marker densities required to represent all haplotype segments within discovery populations (Yan *et al.*, 2009). Linkage disequilibrium among distant loci on the same chromosome or loci on different chromosomes may also occur because of subpopulation structure, kinship, inbreeding, directional selection, and epistasis (Gaut & Long, 2003). Estimating LD between distant loci on the same chromosome or on different chromosomes also is useful for detecting the possibility of false positive associations (Platt *et al.*, 2010). Within outcrossing populations of conifers with large effective population sizes, R^2^ decays rapidly to less than 0.1 over 500 -1500 bases (Brown *et al.*, 2004; Neale & Savolainen, 2004; Pavy *et al.*, 2012a). However, within multi-family pedigrees used for genetic association and genomic selection studies or biparental crosses used for QTL mapping, R^2^ is expected to decay over greater distances proportional to levels of relatedness (Flint-Garcia *et al.*, 2003).

Recently, two dense genetic maps have been published for *P. taeda* consisting of 2466 and 2841 markers that were discovered primarily within genes (Martínez-García *et al.*, 2013; Neves *et al.*, 2014). Neves *et al.* (2014) detected large inversions on four linkage groups relative to a previously published map (Eckert *et al.*, 2010a). Considering that strong co-linearity of markers has consistently been observed between *P. taeda* and other conifer species (Brown *et al.*, 2001; Krutovsky *et al.*, 2004; Pavy *et al.*, 2012b), these inversions were likely attributable to genotyping or mapping errors instead of cytological rearrangements. However, because only two genetic maps were compared, the source of the errors was unclear.

In the present study, a consensus genetic map was constructed by merging three previously published genetic maps for *P. taeda* (Echt *et al.*, 2011; Martínez-García *et al.*, 2013; Neves *et al.*, 2014) with a linkage map from a pseudo-backcross between *P. elliottii* and *P. taeda* (Westbrook *et al.*, 2014). The consensus map positioned 4981 markers via genotyping 1251 individuals from three full-sib populations and one haploid population. Improved functional annotations of the mapped genes were obtained by aligning the partial length expressed sequence tags (ESTs) used for marker discovery against the latest predicted *P. taeda* transcript sequences (NCBI BioProject PRJNA174450; Wegrzyn *et al.*, 2014). Sequences containing mapped markers were also aligned to genomic scaffolds further aiding genome assembly (Zimin *et al.*, 2014). The consensus map was used to compare the genome-wide extent of LD in three distinct types of genotype-phenotype discovery populations before and after accounting for subpopulation structure and kinship. Linkage disequilibrium was compared in ADEPT2 – a population composed of unrelated individuals that has been used for association genetic studies (Quesada *et al.*, 2010; Eckert *et al.*, 2010b; Cumbie *et al.*, 2011), CCLONES – a complex multi-family population that has been used for genomic selection studies (Resende *et al.*, 2012; Westbrook *et al.*, 2013; Westbrook *et al.*, 2014), and BC1 – a single full-sib family derived from a pseudo-backcross between *P. taeda* and *P. elliottii* that has been used for QTL mapping (Muñoz *et al.*, 2011; Westbrook *et al.*, 2014).

## MATERIALS AND METHODS

### Linkage Maps Used to Construct the Consensus Map

Our consensus map was constructed by merging the *P. taeda* 10-5 family map (Neves *et al.*, 2014) with two composite maps from the QTL and BASE pedigrees (Echt *et al.*, 2011; Martínez-García *et al.*, 2013), and the BC1 map from a (*P. taeda* × *P. elliotii* var. *elliottii*) × *P. elliotii* var. *elliottii* pseudo-backcross (Table 1). The 10-5 map was constructed from exome resequencing of 72 haploid megagametophytes from a single tree and contained 2841 SNPs, multiple nucleotide polymorphisms, indels, and presence/absence variants (PAVs) (Neves *et al.*, 2014). The first QTL-BASE map (qtl-base1) contained 460 SSR, RFLP, and ESTP markers genotyped in single F2 cohorts of the QTL and BASE pedigrees (Echt *et al.*, 2011). The second QTL-BASE map (qtl-base2) contained 2466 SNPs, RFLPs, and ESTP markers and was constructed from genotyping two F2 full-sib cohorts in each of the BASE and QTL pedigrees (Martínez-García *et al.*, 2013). All input maps were constructed in JoinMap from three rounds of linkage mapping. Linkage group (LG) numbers and orientation in the input maps were modified to match the historical designations in Echt *et al.* (2011).

**TABLE 1.**
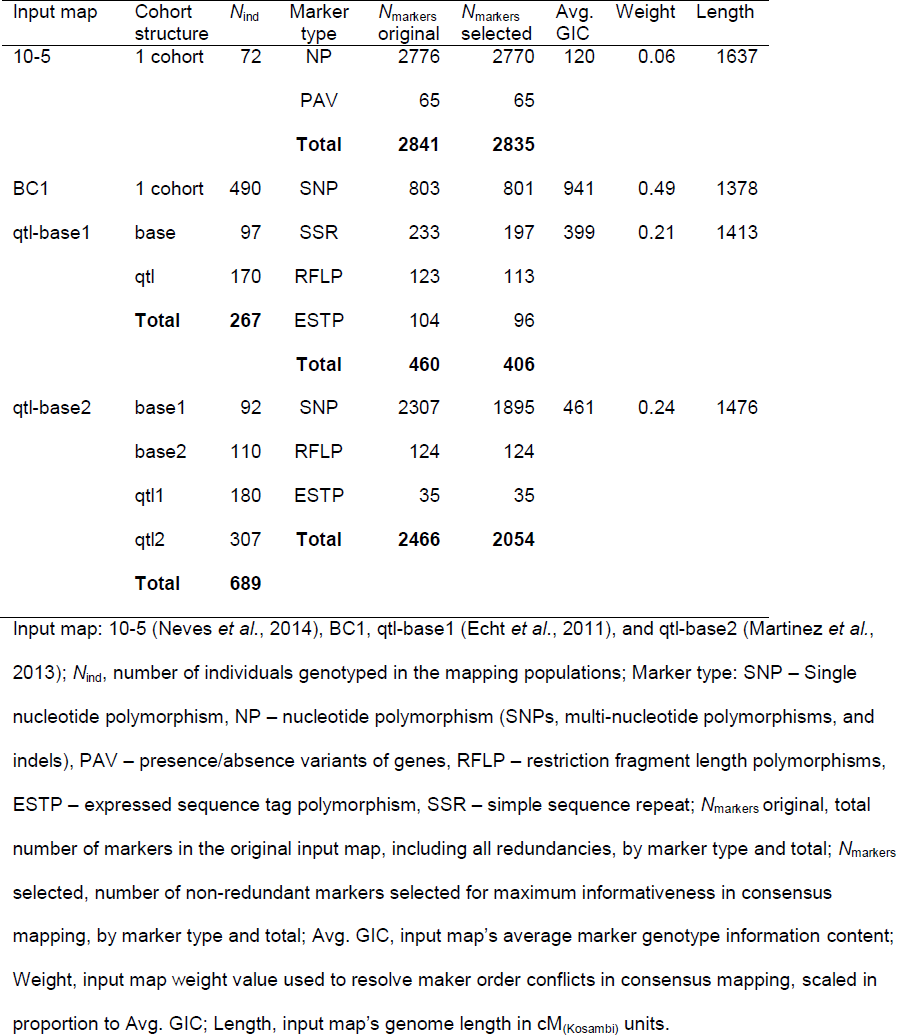
Composition of input maps used to construct *Pinus taeda* consensus maps.

| Input map | Cohort structure | $N_{\text{ind}}$ | Marker type | $N_{\text{markers}}$ original | $N_{\text{markers}}$ selected | Avg. GIC | Weight | Length |
| --- | --- | --- | --- | --- | --- | --- | --- | --- |
| 10-5 | 1 cohort | 72 | NP | 2776 | 2770 | 120 | 0.06 | 1637 |
|  |  |  | PAV | 65 | 65 |  |  |  |
|  |  |  | **Total** | **2841** | **2835** |  |  |  |
| BC1 | 1 cohort | 490 | SNP | 803 | 801 | 941 | 0.49 | 1378 |
| qtl-base1 | base | 97 | SSR | 233 | 197 | 399 | 0.21 | 1413 |
|  | qtl | 170 | RFLP | 123 | 113 |  |  |  |
|  | **Total** | **267** | ESTP | 104 | 96 |  |  |  |
|  |  |  | **Total** | **460** | **406** |  |  |  |
| qtl-base2 | base1 | 92 | SNP | 2307 | 1895 | 461 | 0.24 | 1476 |
|  | base2 | 110 | RFLP | 124 | 124 |  |  |  |
|  | qtl1 | 180 | ESTP | 35 | 35 |  |  |  |
|  | qtl2 | 307 | **Total** | **2466** | **2054** |  |  |  |
|  | **Total** | **689** |  |  |  |  |  |  |

Input map: 10-5 (Neves *et al.*, 2014), BC1, qtl-base1 (Echt *et al.*, 2011), and qtl-base2 (Martinez *et al.*, 2013); $N_{\text{ind}}$ , number of individuals genotyped in the mapping populations; Marker type: SNP – Single nucleotide polymorphism, NP – nucleotide polymorphism (SNPs, multi-nucleotide polymorphisms, and indels), PAV – presence/absence variants of genes, RFLP – restriction fragment length polymorphisms, ESTP – expressed sequence tag polymorphism, SSR – simple sequence repeat; $N_{\text{markers}}$ original, total number of markers in the original input map, including all redundancies, by marker type and total; $N_{\text{markers}}$ selected, number of non-redundant markers selected for maximum informativeness in consensus mapping, by marker type and total; Avg. GIC, input map's average marker genotype information content; Weight, input map weight value used to resolve maker order conflicts in consensus mapping, scaled in proportion to Avg. GIC; Length, input map's genome length in cM<sub>(Kosambi)</sub> units.

### Construction of the BC1 Linkage Map

Previously, a composite map of the BC1 and 10-5 populations was presented in Westbrook *et al.* (2014). Due to large differences in progeny sizes between these populations (490 diploid individuals in BC1 versus 72 haploid megagametophytes in 10-5), we reconstructed a genetic map of the BC1 population separately from the 10-5 map (Neves *et al.*, 2014) prior to its integration into the current consensus map. The BC1 pseudo-backcross population originated from controlled pollination of a *P. taeda* × *P. elliottii* var. *elliottii* F1 hybrid with pollen from a second *P. elliotti* var. *elliottii* individual (Muñoz *et al.*, 2011). Full-sib BC1 progeny, their parents, and the maternal *P. elliottii* grandmother were genotyped at 4861 SNP loci within expressed genes with an Illumina Infinium assay designed for *P. taeda* (Eckert *et al.*, 2010a). Loci that were monoallelic, missing parental genotypes, displayed significant segregation distortion at P < 0.001, or had more than 5% genotyping error rate as inferred from parental genotypes were discarded. For loci that contained genotype errors in less than 5% of the individuals, the erroneous genotypes were recoded as missing data. For genes containing more than one SNP, the marker with the highest genotype information content (described below) was selected for mapping. The BC1 map containing 803 SNPs was constructed in JoinMap v. 4.1 (Van Ooijen, 2011) with three rounds of multi-point regression mapping by specifying the cross-pollinated (CP) population type, a LOD score threshold more than 6 for linkage groups, and the Kosambi mapping function.

### Marker Selection for the Consensus Map

The qtl-base2, BC1, and 10-5 maps were merged based on expressed sequence tag (EST) IDs after omitting the nucleotide position of the SNP within the EST. Nucleotide positions in the 10-5 map did not correspond to the positions in the qtl-base2 and BC1 maps because the SNPs were discovered in different populations and were based upon different alignments. To merge the BC1 and 10-5 maps, which contained one SNP per EST, to the qtl-base2 consensus map, which contained one to three SNPs per EST, it was necessary to select the most informative SNP marker within ESTs from the qtl-base2 map.

Marker informativeness was measured with genotype information content (GIC) calculated as the effective number of genotypic classes (Ge) times the number of individuals genotyped (N_i_). The effective number of genotypic classes for each marker was calculated as 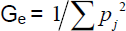 where 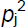 is the squared proportion of the *j*^th^ genotypic class. For biparentally heterozygous (hk × hk) loci, which segregate 1hh:2hk:1kk, G_e_ was calculated using only the homozygous (hh and kk) genotypic classes, while GIC was calculated using the total number of genotyped progeny. The heterozygous (hk) genotype was excluded from the calculation of G_e_ to maintain an inverse linear proportionality of GIC with segregation distortion *χ*^2^ values. For genes genotyped at more than one SNP marker, only the marker with the maximum GIC summed across population cohorts was retained in the qtl-base2 map. The qtl-base1 map was merged to the other maps via RFLP and ESTP markers shared with the qtl-base2 map (Table S1). Files S1 through S4 contain the input maps with GIC used for marker selection.

### Map Weights and Construction of the Consensus Map

Constructing a consensus map directly from recombination frequencies in JoinMap is computationally time consuming and infeasible for multiple populations and cohorts (Wenzl *et al.*, 2006). Instead, the consensus map was constructed directly from the marker IDs and genetic distances in the input maps using two competing algorithms: MergeMap (Wu *et al.*, 2011; http://www.mergemap.org) and LPmerge (Endelman & Plomion 2014; http://cran.r-project.org/web/packages/LPmerge/). Both algorithms use linear programming approaches to resolve marker order conflicts, although MergeMap does so by removing the minimum number of markers from directed acyclic graphs of the input maps, while LPmerge removes inequality constraints among input maps rather than markers. For both approaches, the resolution of marker order conflicts was informed by weighting input maps in proportion to the average GIC of the markers contained in each map. This weighting method incorporated information on number of genotypic classes, segregation distortion, and number of individuals genotyped. For the qtl-base1 and qtl-base2 maps, which were constructed from two and four cohorts of the BASE and QTL populations, respectively, marker GIC was summed across all cohorts in which the marker occurred prior to averaging across markers. The average GIC for the *i*^th^ input map constructed from *k* cohorts and *j* markers was calculated as follows:

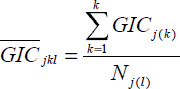

where *GIC*_*j*(*k*)_ is the GIC of the *j*^th^ marker within the *k*^th^ cohort and *N*_*j*(*l*)_ is the number of markers within the *i*^th^ map. Map weights were scaled from 0 to 1 by dividing 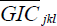 by the sum of the average GIC across maps, 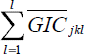.

Root mean squared errors (RMSE) in marker order between both consensus maps and between individual consensus maps and the input maps was calculated with the hydroGOF R package (http://cran.r-project.org/web/packages/hydroGOF/). The consensus map with the overall lowest RMSE with the input maps was used for further analysis.

### Alignment of Mapped Genes to the *P. taeda* Genome and Transcriptome

Expressed sequence tags containing mapped markers were aligned to *P. taeda* genome assembly ver. 1.01 (Neale *et al.*, 2014) using GMAP (Wu & Watanabe, 2005). For ESTs that aligned with more than one genomic scaffold or scaffolds that aligned with two or more ESTs on different LGs, the best alignment was chosen based on alignment length and sequence identity. BlastN (Altschul *et al.*, 1990) 2.2.27+ was used to align ESTs to transcript sequences assembled from RNA-seq reads of the *P. taeda* reference genotype 20-1010 and other genotypes (Mockaitis *et al.*, data available in NCBI BioProject PRJNA174450). Predicted functions of coding sequences were provided from results of Delta-BlastP (Boratyn *et al.*, 2012) alignments of complete transcript protein sequences to the *Arabidopsis thaliana* TAIR10 annotation protein set (Lamesch *et al.*, 2011) and to the NCBI Conserved Domain Database (Marchler-Bauer *et al.*, 2013).

### Genotype-Phenotype Discovery Populations Used for Comparative Linkage Disequilibrium Analysis

The consensus map was used to compare the density of mapped genes and linkage disequilibrium in three genotype-phenotype discovery populations that vary in degree of kinship and subpopulation structure. The discovery populations were 1) BC1, as described above, 2) CCLONES (Comparing Clonal Lines ON Experimental Sites), a population of 923 *P. taeda* progeny in 68 full-sib families generated from circular mating among 54 first and second generation selections from breeding programs in Florida, the Atlantic coastal plain, and the lower gulf states (Baltunis *et al.*, 2007), and 3) ADEPT2 (Allele Discovery of Economic Pine Traits 2), a population consisting of 427 distantly related *P. taeda* individuals sampled from throughout the species range (Eckert *et al.*, 2010a). The CCLONES and ADEPT2 populations were genotyped with an Illumina Infinium assay of 7216 SNP loci (Eckert *et al.*, 2010a). The number of polymorphic loci was 3938 in ADEPT2 and 4854 in CCLONES with 3688 SNPs that were genotyped in both populations.

### Spatial Density of Mapped Genes along Chromosomes

The density of markers along LGs was estimated with kernel density estimation in R (R Core Team, 2012). Fixed bandwidths (*bw*) for the Gaussian kernel density estimator were calculated for each LG following Silverman (1986):

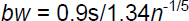

where s is the standard deviation of marker positions in cM (centiMorgan_(Kosambi)_) and *n* is the number of markers per linkage group. Kernel density estimates were multiplied by *n* × *bw* to obtain the number of markers per bandwidth and by *n* to obtain the number of markers per cM. To test for regions where gene density significantly deviated from random expectation, the observed numbers of markers per bandwidth were compared to the 95% confidence interval of a Poisson distribution with mean and variance equal to *n*/(LG length/*bw*). To test for regions where the density of SNPs genotyped in ADEPT2, CCLONES, and BC1 populations differed from random expectations, random selection of equal number of markers from each input population were sampled 1,000 times, without replacement, from the consensus map. Observed marker densities were compared to the 95% confidence interval of marker density derived from 1000 random samples.

### Accounting for Kinship and Structure in the Estimation of Linkage Disequilibrium

The squared correlations (R^2^) between alleles in the CCLONES and ADEPT2 populations were calculated with and without accounting for kinship or subpopulation structure with the R package LDcorSV (Mangin *et al.*, 2012). Kinship in CCLONES was estimated from identity-by-descent proportions expected from pedigree relationships (Henderson, 1976). Subpopulation structure in CCLONES and ADEPT2 populations was inferred with the program fastSTRUCTURE (Raj *et al.*, 2014). In CCLONES, the number of subpopulation clusters (K) tested varied from 2 to 10 using 3037 SNPs, whereas in ADEPT2, K 2 through 7 were tested with 2910 SNPs. Only those SNPs with minor allele frequencies greater than 0.05 were used for structure analysis in both populations. The optimal *K* used to account for population structure in LD analyses was inferred from the model complexity that maximized the marginal likelihood.

## RESULTS

### Comparisons of the Input Maps Used to Construct the Consensus Genetic Map for *Pinus taeda*

Strong concordance in marker order was found between the qtl-base1, qtl-base2, and BC1 maps used to construct the consensus map. By contrast, large inversions were detected in the 10-5 map on LGs 4, 6, 7, and 8 (see Figures S1 – S4). The input maps were merged with 76 to 497 genes that were shared between pairs of maps (see Table S1). Generally, markers shared between maps were located on the same LGs; however, seven markers in the 10-5 map (0_17434, 0_1864, 0_5740, 0_6106, 0_7496, 0_9680, and UMN_213), and two markers in the BC1 map (2_6470 and UMN_1230) were located on different LGs from the qtl-base2 map and were not used in the consensus map.

### Comparisons of the Consensus Maps from Two Map Merging Algorithms

Similar numbers of markers were mapped in the consensus genetic maps generated by MergeMap (4981 markers) and LPmerge (4993 markers) (see Table S2 and Files S5 and S6). The small difference in number of markers in each consensus map was attributable to differences in the number markers removed by the different merging algorithms from a total of 5041 unique markers from the four input maps.

The total length of the MergeMap consensus map (2372 cM) was 1.44 to 1.72 times longer than the lengths of the individual input maps, whereas the length of the LPmerge consensus (1673 cM) was similar to the lengths of the input maps (1378 to 1637 cM) (Table 1). Where there was uncertainty in marker order between the consensus maps, the LPmerge algorithm binned markers into the same map positions, whereas MergeMap assigned unique positions to most markers (see Files S5 and S6). This non-binning attribute of MergeMap accounts for the length expansion of its consensus map compared to the LPmerge map. Comparing the MergeMap and LPmerge consensus maps, small to moderate deviations in marker order were observed for 7 of 12 LGs (see Figure S5). Among these seven LGs, root mean squared error (RMSE) in marker order varied from 7.8 to 23.5 (see Table S3). Larger deviations in marker order between consensus maps were observed for the remaining five LGs. For these five LGs, RMSE values varied from 58.3 to 101.1 (see Figure S5 and Table S3). The MergeMap consensus had lower average RMSE in marker order with the input maps for 11 of 12 LGs as compared to the LPmerge consensus (see Tables S2 and S3). Therefore, the MergeMap consensus map was used for subsequent analyses.

### Summary of the *P. taeda* Consensus Map by Linkage Group

Upon merging the four input maps with MergeMap, portions of LGs that were inverted in the 10-5 map were oriented to the marker order of the other input maps (Figure 1). The MergeMap consensus positioned 4439 SNPs, 197 RFLPs, 197 SSRs, 110 ESTPs, and 63 PAVs (see Files S5 and S7). Between 343 and 501 markers were positioned on individual linkage groups (Table 2). Average and maximum distance between markers was 0.48 cM and 10.9 cM, respectively (Table 2). Linkage group lengths varied from 149.5 cM (LG 12) to 244.4 cM (LG 2). Gaps greater than 10 cM were positioned towards the distal ends of LGs 1, 6, and 7 and originated from the 10-5 and qtl-base2 input maps (see File S7).

**FIGURE 1.**
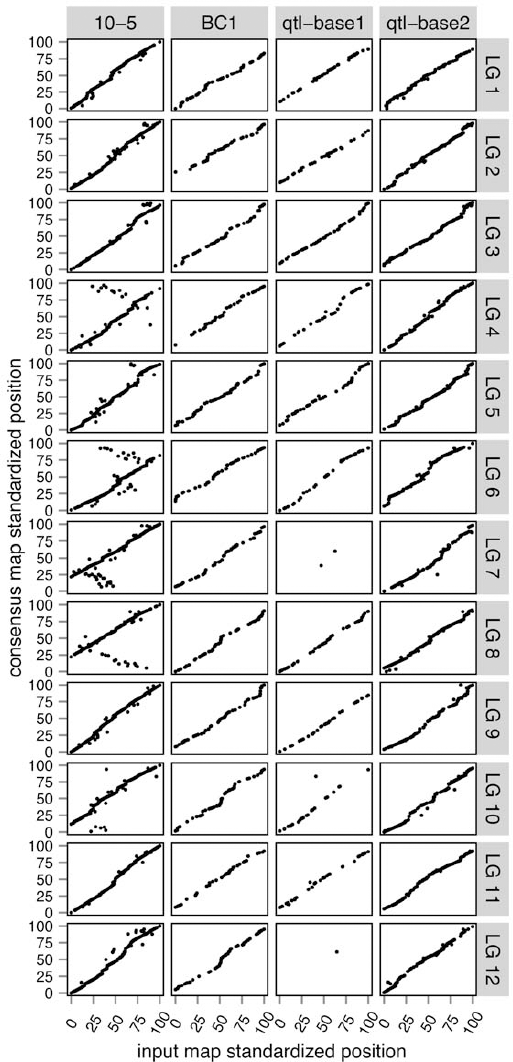
Comparisons of orders of shared markers between input maps and the MergeMap consensus genetic map. Linkage group (LG) lengths were standardized to 100 units for comparison among maps.

**TABLE 2.** Summary of the *Pinus taeda* consensus genetic map by linkage group (LG)

| LG | Length,<br>cM | Average<br>marker<br>spacing, cM | Maximum<br>marker<br>spacing, cM | Observed,<br>*N* markers | Expected,<br>*N* markers | Chi-squared<br>contribution |
| --- | --- | --- | --- | --- | --- | --- |
| 1 | 229.54 | 0.575 | 10.90 | 400 | 482.06 | 13.97 |
| 2 | 244.40 | 0.523 | 5.57 | 468 | 513.27 | 3.99 |
| 3 | 189.01 | 0.470 | 4.66 | 403 | 396.94 | 0.09 |
| 4 | 183.76 | 0.451 | 4.76 | 408 | 385.92 | 1.26 |
| 5 | 207.46 | 0.413 | 3.20 | 503 | 435.69 | 10.40 |
| 6 | 182.92 | 0.418 | 10.55 | 439 | 384.15 | 7.83 |
| 7 | 202.08 | 0.512 | 10.80 | 396 | 424.39 | 1.90 |
| 8 | 203.37 | 0.470 | 6.72 | 434 | 427.1 | 0.11 |
| 9 | 198.48 | 0.489 | 4.39 | 407 | 416.83 | 0.23 |
| 10 | 221.90 | 0.527 | 6.96 | 422 | 466.02 | 4.16 |
| 11 | 149.52 | 0.419 | 5.41 | 358 | 314.01 | 6.16 |
| 12 | 159.33 | 0.465 | 4.02 | 343 | 334.61 | 0.21 |
| Total | **2372** |  |  | **4981** |  | **50.32<sup>a</sup>** |
| Average | 198 | 0.477 | 6.48 | 415 |  |  |

<sup>a</sup> $P = 5.5 \times 10^{-7}$

### Alignment of Mapped Markers to the *P. taeda* Transcriptome and Genome

Of 4981 markers mapped, 4762 (96%), aligned to *P. taeda* transcript assemblies with an average sequence identity of 98.6% (see File S5). Predicted functions for 4715 mapped ESTs were obtained through alignment of the translated transcripts to 2954 unique *Arabidopsis* protein sequences in the TAIR10 database and to 1727 unique conserved domains in the NCBI CDD database (see File S5).

A total of 4881 mapped markers were aligned to 4225 *P. taeda* genomic scaffolds with 98.3% average nucleotide sequence identity (see File S5). Ratios of physical to genetic distance were calculated for 47 pairs of mapped ESTs that aligned to single genomic scaffolds, but to different transcripts with unique annotations. Mean physical to genetic distance among these 47 marker pairs was 301400 bases per cM with a range of 106 to 5867000 bases per cM. These ratios were considerably less than the expectation of 9.15 Mb per cM estimated from a genome size of 21.6 Gb (O’Brien *et al.*, 1996) and a total map length of 2372 cM (Table 2).

### Variation in Marker Density along Chromosomes

The observed number of markers per LG differed significantly from expectation assuming a uniform density of markers per cM(*χ*^2^ _11 df_ = 50.32, *p* = 5.5 × 10^−7^; Table 2). Linkage groups 5, 6, and 11 had substantially more markers than expected, while LGs 1, 2, and 10 had fewer markers than expected. Figure 2a compares observed variation in the number of markers per cM to variation expected from a random Poisson process. Lower than expected marker densities were observed toward the distal ends of all 12 LGs, while greater than expected marker densities were observed in putative centromeric regions of linkage groups 2, 3, 6, 7, and 8 (Figure 2a).

### Mapping Genes in Three Genotype-Phenotype Discovery Populations

The consensus map positioned 2673 of 3938 SNPs (68%) currently genotyped in unrelated individuals from the *P. taeda* ADEPT2 population, 3046 of 4854 SNPs (63%) in the CCLONES multiple-family pedigree, and 999 of 1032 SNPs (97%) among BC1 full-sibs (Table 3). Average distance between adjacent mapped SNPs was approximately one cM in ADEPT2 and CCLONES and 2.5 cM in BC1 (Table 3). Figure 2b-d compares the densities of markers mapped in each population with 1) the marker densities from the consensus map and 2) the 95% confidence interval of densities obtained from randomly sampling number of markers mapped in each population from the consensus map. Marker densities (N cM^−1^) varied from 0.03 to 1.5 in ADEPT2 (mean = 0.89), from 0.04 to 1.63 in CCLONES (mean = 0.97), and from 0.006 to 0.68 in BC1 (mean = 0.36). Marker densities deviated from random expectations in similar genetic map positions in the ADEPT2 and CCLONES populations (Figure 2b, c).

**TABLE 3.**
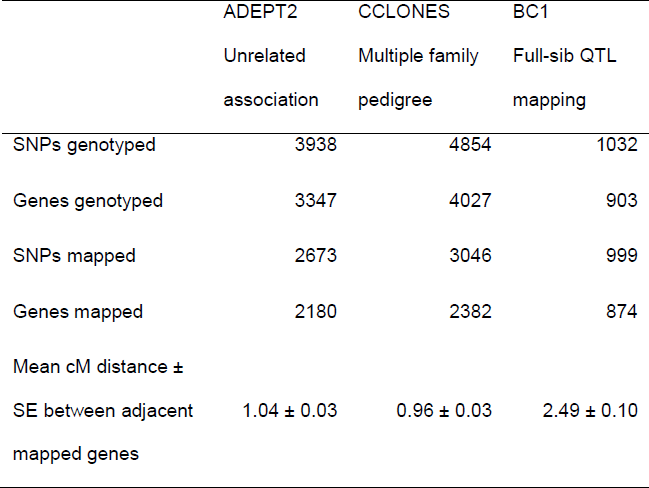
Number of SNPs and genes mapped in three types of genotype-phenotype discovery populations.

**FIGURE 2.**
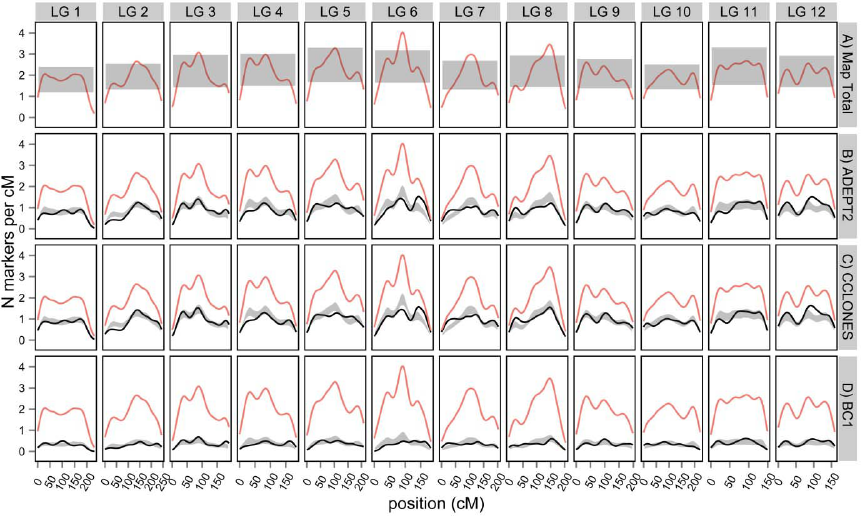
Kernel density estimation of mapped genes in the MergeMap *Pinus taeda* consensus genetic map. A) Estimated marker densities (red lines) compared against the 95% confidence interval of a Poisson distribution (grey regions) with mean and variance (λ) calculated assuming an equal number of markers per bandwidth (λ = Nmarkers/([LG length/bandwidth]). B) through D) estimated densities of mapped polymorphic genes in ADEPT2 (unrelated association), CCLONES (multiple-family pedigree), and BC1 (full-sib QTL mapping) genotype-phenotype discovery populations. The densities of genes mapped in each population (black lines) were compared against gene densities in the consensus map (red lines), and the 95% confidence interval of density obtained from randomly sampling the number of genes mapped in each population from the consensus map 1,000 times.

### Genome-wide Extent of Linkage Disequilibrium

Within the ADEPT2, CCLONES, and BC1 populations, respectively, 575, 805 and 114 genes each had two SNP loci genotyped. The distributions of intragenic R^2^ values were bimodal in CCLONES and ADEPT2, with a high frequency of R^2^ values that were approximately zero and a lower frequency of R^2^ values that were approximately one (Figure 3). By contrast, the modal value of R^2^ between SNPs in the same gene was approximately one among BC1 full-sibs.

**FIGURE 3.**
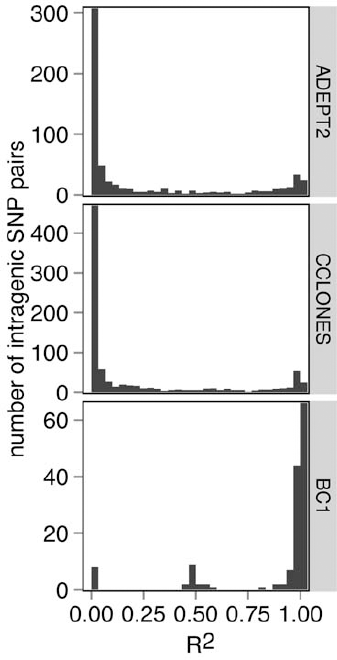
Distributions of squared correlation (R^2^) between pairs of SNPs within the same genes. In the ADEPT2 (unrelated association), CCLONES (multiple-family pedigree), and BC1 (full-sib QTL mapping) populations, respectively, 575, 805, and 114 genes were genotyped at two SNP loci.

Average R^2^ between intergenic markers separated by less than one cM was 0.04 in ADEPT2 and 0.05 in CCLONES. Percentages of SNP pairs less than 1 cM apart with R^2^ greater than 0.1 varied from 2.7% to 9.7% in ADEPT2 and from 4.7% to 12.4% in CCLONES depending on minimum minor allele frequency (MAF) thresholds (Table 4). Average R^2^ did not decay substantially with genetic distance in ADEPT2 and CCLONES (Figure 4). By contrast, strong and extended linkage disequilibrium was observed in BC1. Average R^2^ was 0.8 between SNPs less than one cM apart and R^2^ decayed to less than 0.1 over 53 cM (Figure 4).

**TABLE 4.**
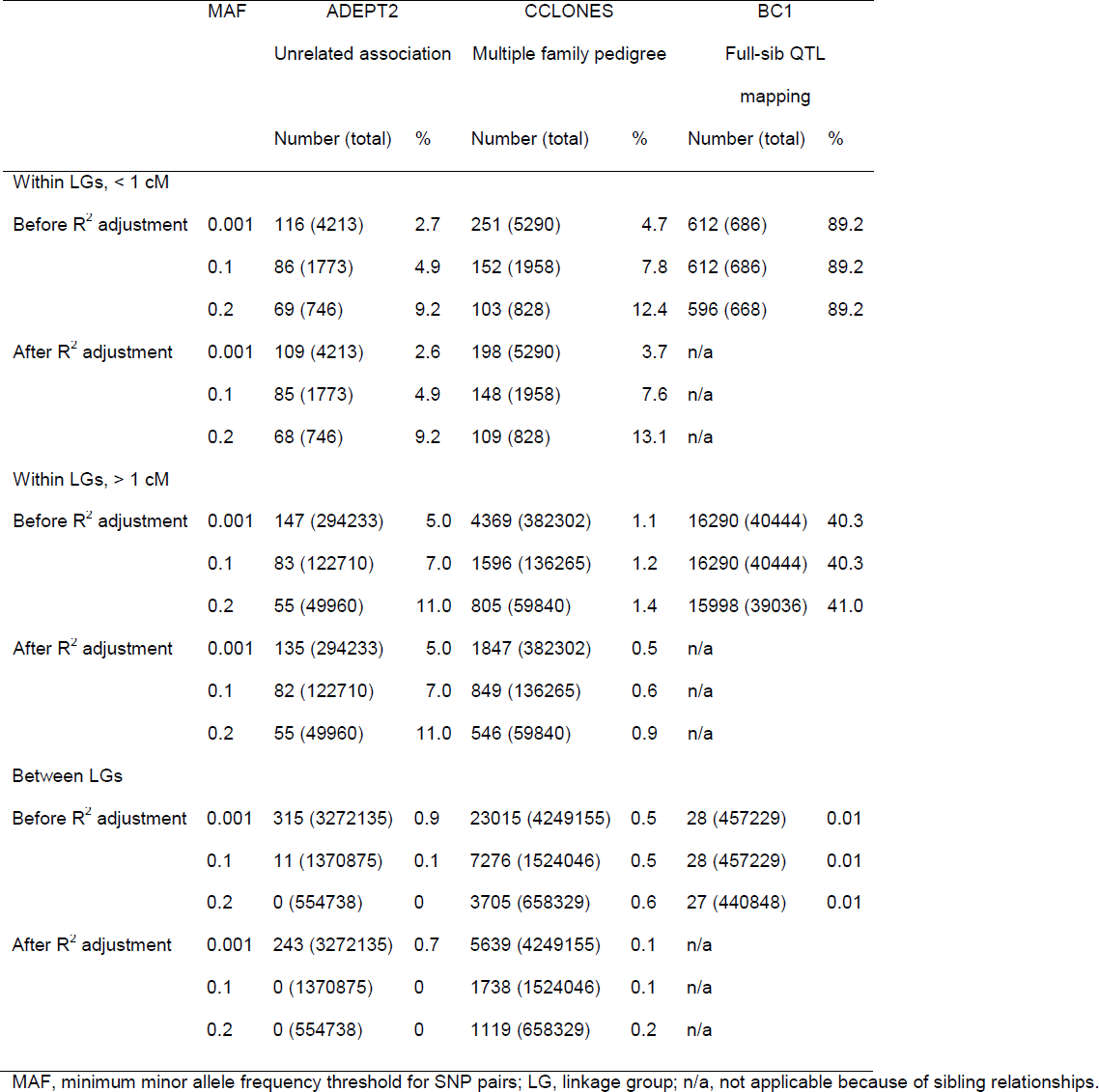
Number and percentage of SNPs pairs among different genes with R^2^ allele correlations > 0.1. R^2^ for three linkage distance classes, each before and after adjusting R^2^ for subpopulation structure in three types of genotype-phenotype discovery populations.

|  | MAF | ADEPT2 |  | CCLONES |  | BC1 |  |
| --- | --- | --- | --- | --- | --- | --- | --- |
|  |  | Unrelated association |  | Multiple family pedigree |  | Full-sib QTL mapping |  |
|  |  | Number (total) | % | Number (total) | % | Number (total) | % |
| Within LGs, < 1 cM |  |  |  |  |  |  |  |
| Before $R^2$ adjustment | 0.001 | 116 (4213) | 2.7 | 251 (5290) | 4.7 | 612 (686) | 89.2 |
|  | 0.1 | 86 (1773) | 4.9 | 152 (1958) | 7.8 | 612 (686) | 89.2 |
|  | 0.2 | 69 (746) | 9.2 | 103 (828) | 12.4 | 596 (668) | 89.2 |
| After $R^2$ adjustment | 0.001 | 109 (4213) | 2.6 | 198 (5290) | 3.7 | n/a | |
|  | 0.1 | 85 (1773) | 4.9 | 148 (1958) | 7.6 | n/a |  |
|  | 0.2 | 68 (746) | 9.2 | 109 (828) | 13.1 | n/a |  |
| Within LGs, > 1 cM |  |  |  |  |  |  |  |
| Before $R^2$ adjustment | 0.001 | 147 (294233) | 5.0 | 4369 (382302) | 1.1 | 16290 (40444) | 40.3 |
|  | 0.1 | 83 (122710) | 7.0 | 1596 (136265) | 1.2 | 16290 (40444) | 40.3 |
|  | 0.2 | 55 (49960) | 11.0 | 805 (59840) | 1.4 | 15998 (39036) | 41.0 |
| After $R^2$ adjustment | 0.001 | 135 (294233) | 5.0 | 1847 (382302) | 0.5 | n/a | |
|  | 0.1 | 82 (122710) | 7.0 | 849 (136265) | 0.6 | n/a |  |
|  | 0.2 | 55 (49960) | 11.0 | 546 (59840) | 0.9 | n/a |  |
| Between LGs |  |  |  |  |  |  |  |
| Before $R^2$ adjustment | 0.001 | 315 (3272135) | 0.9 | 23015 (4249155) | 0.5 | 28 (457229) | 0.01 |
|  | 0.1 | 11 (1370875) | 0.1 | 7276 (1524046) | 0.5 | 28 (457229) | 0.01 |
|  | 0.2 | 0 (554738) | 0 | 3705 (658329) | 0.6 | 27 (440848) | 0.01 |
| After $R^2$ adjustment | 0.001 | 243 (3272135) | 0.7 | 5639 (4249155) | 0.1 | n/a | |
|  | 0.1 | 0 (1370875) | 0 | 1738 (1524046) | 0.1 | n/a |  |
|  | 0.2 | 0 (554738) | 0 | 1119 (658329) | 0.2 | n/a |  |

MAF, minimum minor allele frequency threshold for SNP pairs; LG, linkage group; n/a, not applicable because of sibling relationships.

**FIGURE 4.**
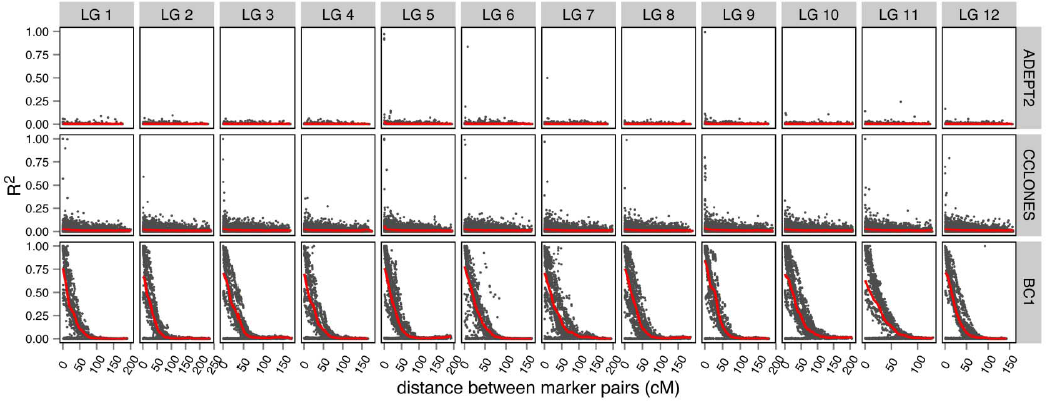
Squared correlation (R^2^) between alleles at different loci versus genetic distance in ADEPT2 (unrelated association), CCLONES (multiple-family pedigree), and BC1 (full-sib QTL mapping) discovery populations. Displayed are values of R^2^ between mapped SNPs in different ESTs with minor allele frequencies greater than 0.1 and with less than 50% missing data. Kernel regression of mean R^2^ versus genetic distance (red lines) was carried out with the ksmooth function in R.

Extended linkage disequilibrium, defined as SNP pairs more than one cM apart with R^2^ values greater than 0.1, was rare in ADEPT2, occurring between only 0.05% of SNP pairs (Table 4). Extended LD was more prevalent in CCLONES, occurring between 1.1% of SNP pairs greater than one cM apart. Furthermore, the range of genetic distances over which extended linkage disequilibrium was observed was greater in CCLONES as compared to ADEPT2 (Figure 5a, Table 4). Extended LD was observed between SNPs more than 100 cM apart in CCLONES, whereas LD rarely extended more than 50 cM in ADEPT2 (Figure 5a). Linkage disequilibrium between SNPs on different LGs was also more prevalent in CCLONES than in ADEPT2 or BC1 (Table 5; Figure 6). In CCLONES, 0.54% SNP pairs on different LGs had R^2^ > 0.1. Smaller percentages of SNPs on different LGs had R^2^ > 0.1 in ADEPT2 (0.009%) and BC1 (0.006%) (Table 4).

**FIGURE 5.**
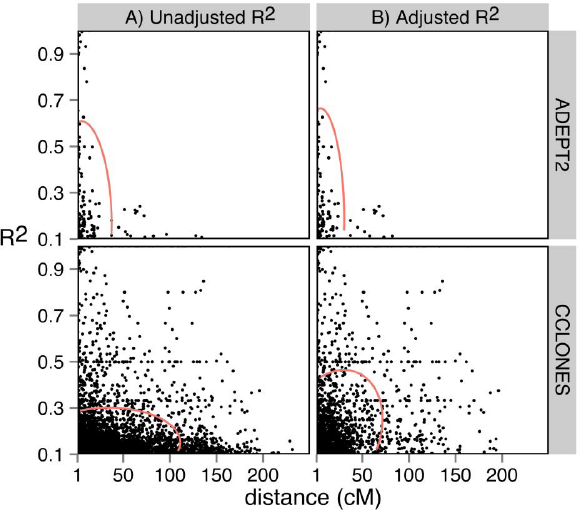
Extended linkage disequilibrium within linkage groups in the ADEPT2 (unrelated association) and CCLONES (multiple-family pedigree) populations. Values of R^2^ versus genetic distance are plotted for SNP pairs with R^2^ > 0.1 that were more than one cM apart. Values of R^2^ were estimated A) before and B) after accounting for population structure in ADEPT2 and kinship in CCLONES with the R package LDcorSV (Mangin *et al.*, 2012). The 95% confidence ellipses of R^2^ versus genetic distance are depicted in red.

**FIGURE 6.**
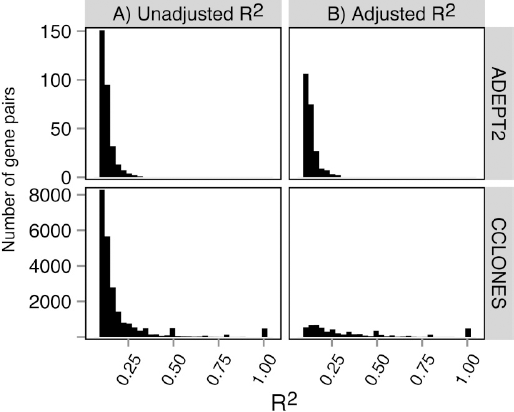
Linkage disequilibrium between genes on different linkage groups in the ADEPT2 and CCLONES populations. Distributions R^2^ were plotted for SNP pairs on different LGs with R^2^ > 0.1. The distributions were compared A) before and B) after adjusting R^2^ values for subpopulation structure in ADEPT2 and kinship in CCLONES.

### The Effects of Subpopulation Structure and Kinship on Extended Linkage Disequilibrium

Squared correlations (R^2^) between pairs of SNP genotypes were compared before and after accounting for subpopulation structure in ADEPT2 and kinship in CCLONES to quantify the effects of these factors on the observed patterns of LD (Figures 5 and 6). In ADEPT2, a model with two subpopulations had the greatest marginal likelihood in fastSTRUCTURE analysis (see Figure S6a); however, the K=2 structure matrix was not invertible and could not be utilized for linkage disequilibrium analyses. Therefore, the structure matrix with three subpopulations (see File S8), the second most likely K, was utilized to account for structure in the analysis of LD in ADEPT2.

Kinship, as measured by identity-by-descent (IBD) between individuals, was expected to be zero among distantly related individuals in ADEPT2. In CCLONES, a continuous increase in marginal likelihood from one to ten subpopulations was observed in fastSTRUCTURE analysis indicating that subpopulation structure was weak or non-existent (see Figure S6b). Identity-by-descent proportions among pairs of individuals in the CCLONES pedigree varied from 0 to 0.5; however, average IBD between all individuals was small (0.043) due to the fact that 80% of individuals were unrelated (see Table S4). Values of R^2^ were not adjusted for kinship or structure in BC1 because expected IBD proportions were uniformly 0.5 among full-sibs and no subpopulation structure was expected from a bi-parental cross.

In ADEPT2, of the 147 SNP pairs in extended LD within LGs, 135 pairs also had R^2^ values that were greater than 0.1 after accounting for subpopulation structure (Table 4). Between LGs, accounting for structure reduced the number of SNP pairs with R^2^ greater than 0.1 from 315 to 243. Thus, accounting for structure in ADEPT2 resulted in an 8% reduction in the number of SNP pairs in extended LD within LGs and a 23% reduction between LGs.

Larger reductions in LD were observed after accounting for kinship in CCLONES. The number of SNPs in extended LD was reduced by 47% within LGs and 75% between LGs (Table 4; Figure 6). Furthermore, accounting for kinship reduced the average genetic distances between SNPs in extended LD (Figure 5b). No SNP pairs in extended LD within LGs and one pair in LD between LGs were shared between ADEPT2 and CCLONES after accounting for structure or kinship, respectively (see File S9).

## DISCUSSION

Nearly 5,000 markers, primarily located within expressed transcripts, were positioned within the *P. taeda* consensus genetic map. The consensus map contained 1.8 to 10.8 times the number of markers of the input maps (Table 1) and genetically mapped approximately 10% of the 50,172 genes predicted for *P. taeda* (Neale *et al.*, 2014). The strong co-linearity in gene order between the qtl-base1, qtl-base2, and BC1 maps, in contrast with the large inversions on four linkage groups in the 10-5 map, indicates that inversions in the 10-5 map were most likely mapping errors (Figure 1, also see Figures S1 – S4). This interpretation is supported by the strong synteny of homologous markers among conifer genetic maps (Brown *et al.*, 2001; Krutovsky *et al.*, 2004; Pavy *et al.*, 2012b) and the fact that the 10-5 map was constructed from a much smaller population of meiotic products of 72 haploid megagametophytes (Table 1), which would have decreased the probability of correct marker ordering relative to the larger mapping populations (Maliepaard *et al.* 1997).

Alignment of genetically mapped sequences to the *P. taeda* genome enabled the estimation of physical to genetic distances for a small number of marker pairs that aligned to single genomic scaffolds. There are two reasons why the mean of 301400 bases per cM probably represents a lower bound for physical to genetic distances in the *P. taeda* genome. First, it is possible that some distance estimates were erroneous due to paralogous genes incorrectly aligning to the same genomic scaffold, although we attempted to exclude this possibility by considering only marker pairs that aligned to different transcripts with unique annotations. Second and more compellingly, given an average marker spacing of 0.48 cM (Table 2) and an expectation of 9.15 Mb per cM, scaffold lengths would have to be at least 4.39 Mb to align to two averagely spaced markers. The mean length of scaffolds aligned to mapped markers, however, was 318 kb and the maximum length was 5.77 Mb (see File S5). Therefore, the majority of scaffolds were too short to span multiple markers resulting in an artificially low mean physical to genetic distance. Integration of the consensus genetic map with a physical map can aid in genome assembly and enable genome wide estimates of recombination rates, as has been done for other plant species (*e.g.*, Chen *et al.*, 2002).

The density of mapped markers varied from less than one per cM toward the distal ends of LGs to four per cM in more gene rich regions towards the middle of LGs (Figure 2). Similar patterns of mapped gene densities have been observed in the consensus genetic maps of *Picea glauca* (white spruce) and *Picea mariana* (black spruce) and *Pinus pinaster* (maritime pine) (Pavy *et al.*, 2012; Plomion *et al.*, 2014). Gene rich regions appear to occur in centromeric regions with reduced recombination rates, while gene poor regions occur near telomeres with higher rates of recombination. Power to detect associations between markers and traits is greater in gene rich regions with reduced recombination rates, but the genomic resolution to fine-map casual variants in these regions is reduced (Nachman, 2002).

### Comparison of LD Extent between Discovery Populations: Implications for Comparative QTL and Association Mapping across Populations

Compared to the largest previously published map for *P. taeda*, a composite map of the BC1 and 10-5 populations with 3353 genes (Westbrook *et al.*, 2014), the consensus map presented here increased the number polymorphic genes mapped from 1236 to 2180 (+76%) in ADEPT2, from 1373 to 2382 (+73%) in CCLONES, and from 803 to 874 in BC1 (+9%). The extent of LD was expected to be high in the BC1 full-sib population and low between unrelated individuals in ADEPT2. Expectations for the extent of LD were uncertain in CCLONES, which is composed primarily of full-sibs, half-sibs, and unrelated individuals (see Table S4). In CCLONES, LD tended to be weak between SNPs within genes (Figure 3), in different genes on the same LG (Figure 4), and on different LGs (Table 4). Patterns of LD in CCLONES more closely resembled those of unrelated individuals in ADEPT2 than the strong and extended LD observed among BC1 full-sibs (Figures 3 and 4). Low levels of linkage disequilibrium in CCLONES may be explained by the fact that 80% of pairs of individuals within the pedigree were unrelated (see Table S4). Each of the 54 parents from the CCLONES pedigree were mated to only two to six other parents (average 4.3); therefore, many individuals did not share common parents (Baltunis *et al.*, 2007; Munoz *et al.*, 2014).

In ADEPT2 and CCLONES, low average R^2^ values between genes on the same LG (R^2^_avg_ < 0.05) and the lack of substantial decay of R^2^ with genetic distance precluded the estimation marker densities required to represent non-recombined haplotype segments of LGs (Figure 4). This result implies that association genetic studies in these populations are underpowered to comprehensively detect casual variants at current marker densities. This is not surprising considering that LD decays within hundreds to thousands of bases in outcrossing *P. taeda* populations (Brown *et al.*, 2004; Neale & Savolainen, 2004). Furthermore, only 4027 genes in CCLONES and 3347 genes in ADEPT2 were genotyped at polymorphic SNP loci (Table 3) of an estimated 50000 genes predicted in the *P. taeda* genome (Neale *et al.*, 2014).

Although LD tended to be weak in CCLONES and ADEPT2, rare cases of extended LD were observed within and between LGs (Table 4). The number of SNP pairs in extended LD and the genetic distances between them was greater in CCLONES as compared to ADEPT2 (Table 4; Figure 5). Furthermore, greater reductions in the number and distance between SNPs in extended LD were observed after accounting for kinship in CCLONES versus structure in ADEPT2 (Table 4). The relatively small effects of structure on patterns of LD may be explained by high rates of gene flow and weak subpopulation structure across the geographic range of *P. taeda* (Al’Rabab’ah & Williams, 2002; Eckert *et al.*, 2010a; Chhatre *et al.*, 2013). The larger effect of kinship on LD underscores the importance of accounting for kinship in association genetic studies to reduce the probability of false positives arising from extended LD.

Small percentages of SNP pairs remained in extended LD after accounting for structure or kinship (Table 4). Accounting for these factors reduces bias in mean R^2^ values (Mangin *et al.*, 2012), but may only partially adjust individual observations of R^2^. Extended LD within LGs may also occur as the result of directional selection or recombination cold spots (Nachman, 2002; Gaut & Long, 2003). Regardless of the mechanisms underlying rare cases of extended LD, SNPs in LD after structure or kinship correction may be queried to detect potential for false positive associations (see File S9).

The consensus map and LD analysis enables comparison of locations of genetic associations and QTL for traits phenotyped in multiple populations. For example, Westbrook *et al.*, (2014) used a composite map of the 10-5 and BC1 populations to compare the locations of SNPs associated with resin canal number in CCLONES to QTL intervals of same trait in BC1. The point locations of associated SNPs could also be compared between populations (e.g., CCLONES and ADEPT2) if the associated SNPs are in LD within at least one the populations (Teo *et al.*, 2010).

### Conclusions

The consensus genetic map for *P. taeda* presented here is, to our knowledge, the most densely populated linkage map for a conifer to date (Ritland *et al*, 2011). Improved functional annotations of mapped genes via alignment with full-length transcripts will be useful for the discovery of candidate genes underlying traits and for comparisons of gene order between *P. taeda* and other species (Pavy *et al.*, 2012). Furthermore, the consensus map coupled with the genome-wide analysis of linkage disequilibrium in three discovery populations establishes a foundation for comparative association and QTL mapping and the implementation of genomic selection in pine.

## ACKNOWLEDGEMENTS

JWW was supported by a USDA CSREES Food and Agricultural Sciences National Needs Graduate Fellowship, VEC was supported by USDA NIFA Award #2011-68002-30185 (PINEMAP) and the USDA Forest Service. LSW was supported by the National Science Foundation under Grant No. ABI-1062432 to Indiana University. PMG, DN, and KM were supported in part by USDA NIFA Award #2011-67009-30030 (PineRefSeq) to University of California, Davis.

## Supporting Information

**FIGURE S1.**
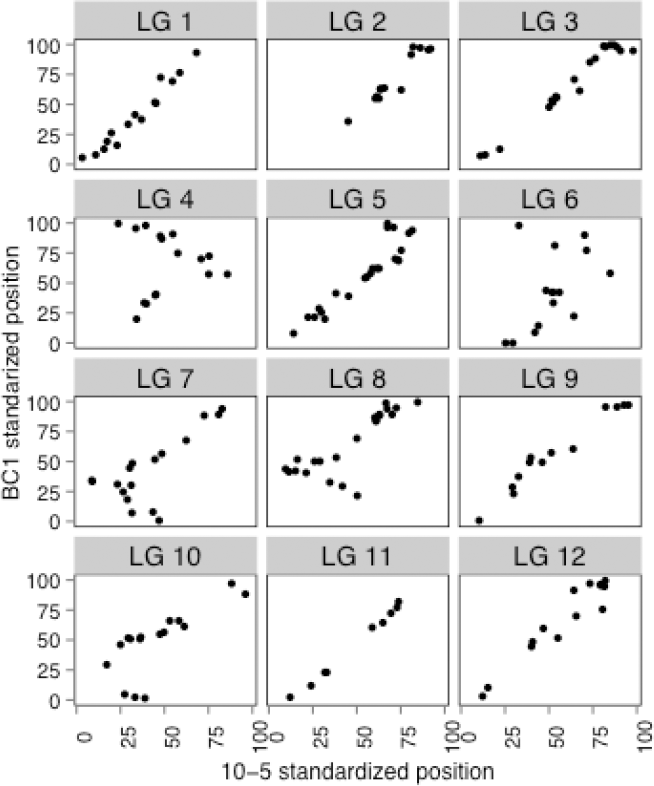
Comparison of order of shared markers between the 10-5 and BC1 input maps; linkage group lengths were standardized to 100 units for comparison between maps.

**FIGURE S2.**
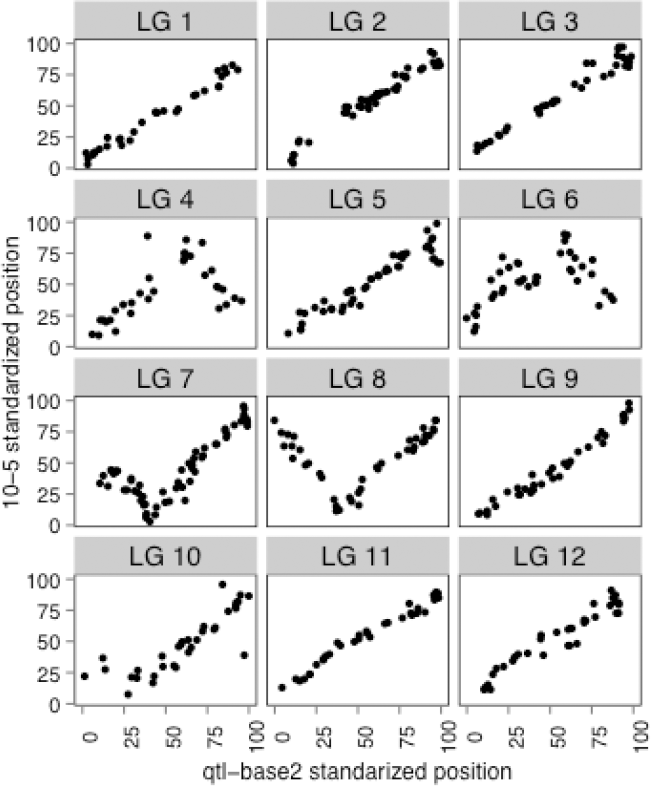
Comparison of order of shared markers between the qtl-base2 and 10-5 input maps

**FIGURE S3.**
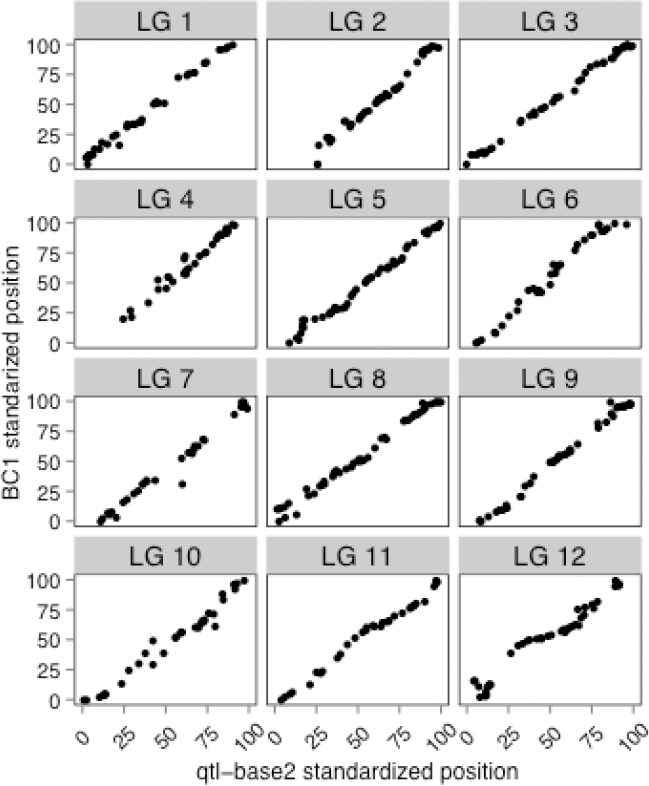
Comparison of order of shared markers between the qtl-base2 and BC1 input maps

**FIGURE S4.**
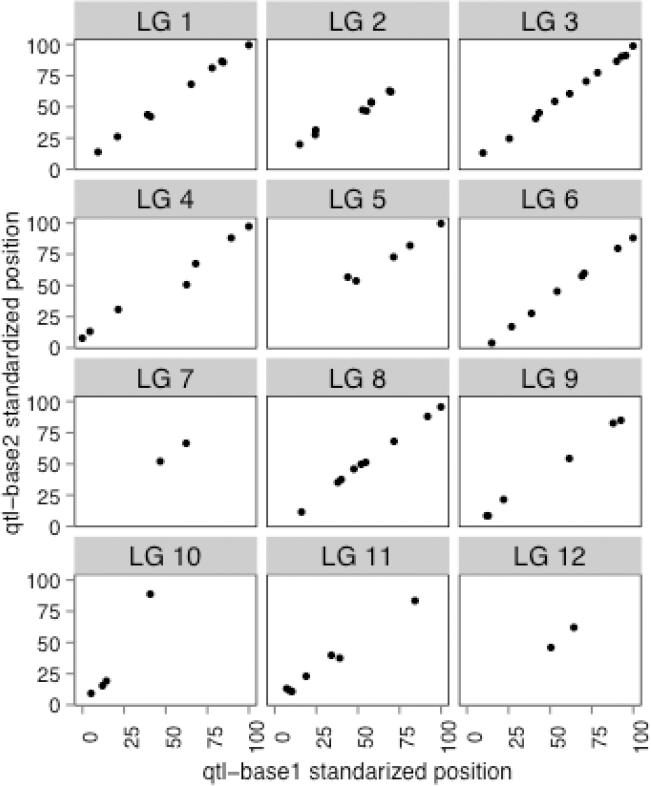
Comparison of order of shared markers between the qtl-base1 and qtl-base2 input maps

**FIGURE S5.**
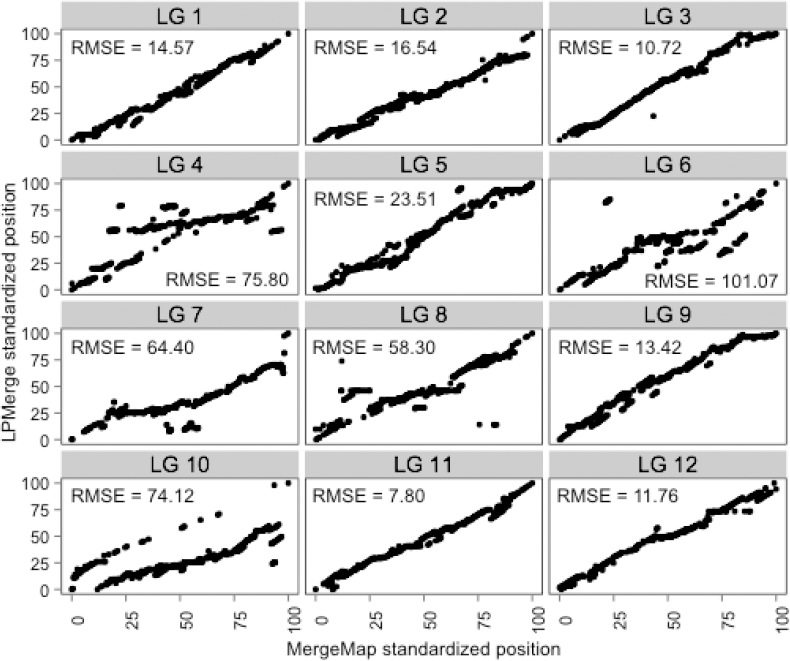
Comparison of order of shared markers between the MergeMap and LPMerge consensus genetic maps for *P. taeda*; Root mean squared error (RMSE) in marker between the consensus maps was estimated for each linkage group.

**FIGURE S6.**
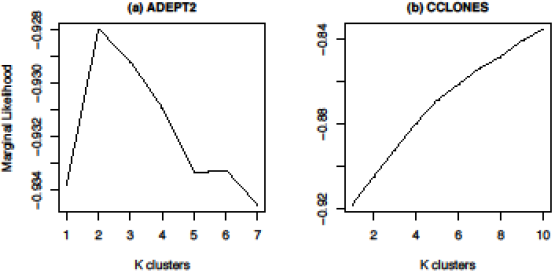
Population structure analysis in ADEPT2 (a) and CCLONES (b) populations with fastSTRUCTURE (Raj *et al.* 2013) marginal likelihood over a range of K values

**TABLE S1.** Number of markers included and shared among input maps used to construct the *P. taeda* MergeMap consensus genetic map; the diagonal represents the number of markers selected for inclusion in the consensus map from each input map, while the off-diagonal elements represent the number of markers shared between maps.

|  | 10-5 | BC1 | qtl-base1 | qtl-base2 |
| --- | --- | --- | --- | --- |
| 10-5 | 2835 |  |  |  |
| BC1 | 152 | 801 |  |  |
| qtl-base1 | 0 | 0 | 406 |  |
| qtl-base2 | 477 | 497 | 76 | 2054 |

**TABLE S2.** Comparisons of the MergeMap and LPmerge P. taeda consensus genetic maps.

| Map attribute | MergeMap | LPmerge |
| --- | --- | --- |
| Number of markers mapped | 4981 | 4993 |
| Number of unique marker positions | 4916 | 3568 |
| Average marker spacing (cM) | 0.477 | 0.356 |
| Total map length (cM) | 2371.77 | 1673.09 |
| RMSE with input maps averaged across LGs | 4.98 | 10.93 |

**TABLE S3.** Root mean squared error (RMSE) in marker order between the MergeMap or LPmerge consensus genetic maps and the input maps, by linkage group (LG).

| LG | 10-5 | BC1 | qtl-base1 | qtl-base2 | Mean RMSE | Std dev. RMSE |
| --- | --- | --- | --- | --- | --- | --- |
| **MergeMap** |  |  |  |  |  |  |
| 1 | 4.11 | 0 | 2.41 | 0.3 | 1.71 | 1.93 |
| 2 | 6.33 | 0 | 1.57 | 1.16 | 2.27 | 2.79 |
| 3 | 4.64 | 0 | 2.67 | 0 | 1.83 | 2.26 |
| 4 | 27.96 | 0.16 | 2.3 | 0.77 | 7.8 | 13.47 |
| 5 | 11.63 | 0 | 1.48 | 1.47 | 3.64 | 5.37 |
| 6 | 32.68 | 0 | 3.04 | 0 | 8.93 | 15.9 |
| 7 | 29.84 | 0 | 4.11 | NA | 11.32 | 16.17 |
| 8 | 36.81 | 0 | 3.31 | 0 | 10.03 | 17.92 |
| 9 | 5.55 | 0 | 3.46 | 0 | 2.25 | 2.74 |
| 10 | 16.37 | 0 | 2.33 | 1.75 | 5.11 | 7.57 |
| 11 | 2.13 | 0 | 1.14 | 1.04 | 1.08 | 0.87 |
| 12 | 8.06 | 0 | 3.4 | NA | 3.82 | 4.05 |
| **LPmerge** |  |  |  |  |  |  |
| 1 | 8.93 | 1 | 4.55 | 0 | 3.62 | 4.04 |
| 2 | 11.74 | 0 | 3.76 | 0 | 3.87 | 5.54 |
| 3 | 7.29 | 2.14 | 4.69 | 0 | 3.53 | 3.16 |
| 4 | 47.99 | 12.59 | 25.25 | 0 | 21.46 | 20.47 |
| 5 | 18.58 | 1.74 | 3.74 | 0 | 6.02 | 8.52 |
| 6 | 75.73 | 12.01 | 29.77 | 0 | 29.38 | 33.23 |
| 7 | 55.9 | 2.88 | 11.01 |  | 23.26 | 28.55 |
| 8 | 42.24 | 6.57 | 21.86 | 0 | 17.67 | 18.77 |
| 9 | 8.96 | 0.5 | 4.9 | 0 | 3.59 | 4.2 |
| 10 | 16.46 | 9.53 | 27.44 | 0 | 13.36 | 11.56 |
| 11 | 4.2 | 0 | 2.94 | 0 | 1.78 | 2.12 |
| 12 | 8.93 | 1 | 4.55 | 0 | 3.62 | 4.04 |

**TABLE S4.** Summary of identity by descent (IBD) proportions among pairs of individuals in the CCLONES pedigree.

| IBD proportion | N pairs | % of pedigree |
| --- | --- | --- |
| 0 | 338676 | 79.59 |
| 0.0625 | 15545 | 3.65 |
| 0.125 | 18258 | 4.29 |
| 0.1875 | 1123 | 0.26 |
| 0.25 | 43872 | 10.31 |
| 0.3125 | 1090 | 0.26 |
| 0.375 | 527 | 0.12 |
| 0.5 | 6412 | 1.51 |

## Supporting Files

All.txt Supporting Files are comma-delimited data tables with missing data as NA, in Unix text file format with UTF-8 encoding.

File S1 – The 10-5 linkage map with marker GIC (.txt)

File S2 – The BC1 linkage map with marker GIC (.txt)

File S3 – The qtl-base1 linkage map with marker GIC (.txt)

File S4 – The qlt-base2 linkage map with marker GIC (.txt)

File S5 – MergeMap consensus genetic map for *Pinus taeda* with genomic scaffold assignments and transcript and protein annotations (.txt)

File S6 – LPmerge consensus genetic map for *Pinus taeda* (.txt)

File S7 – Mapchart of MergeMap consensus map depicting the 12 linkage groups and 4981 loci graphed at a proportional scale; marker names and notations are readable at 600% zoom. (PDF)

File S8 – fast STRUCTURE results matrix of the ADEPT2 population for three subpopulations, K=3. (.txt)

File S9 – Pairs of expressed sequences containing SNPs in extended LD (R2 >0.1), before and after accounting for structure in ADEPT2 and kinship in CCLONES; table includes MAF and consensus map positions, (.txt)

